# Visual salience of the stop-signal affects neural dynamics of controlled inhibition

**DOI:** 10.1101/299560

**Authors:** Pani Pierpaolo, Giarrocco Franco, Giamundo Margherita, Montanari Roberto, Brunamonti Emiliano, Ferraina Stefano

**Author notes:** equally contributed (Pani Pierpaolo, Giarrocco Franco). Correspondence: Pierpaolo Pani PhD, Department of Physiology and Pharmacology, P.le Aldo Moro 5 - 00185 Rome, Italy. Author Contributions: Designed research: PP,SF,RM; Performed Research: FG,RM,PP,MG,EB; Analyzed data: PP,FG,EB,MG; Wrote the paper: PP,SF,FG.

## Abstract

The countermanding or stop-signal task is broadly used to evaluate response inhibition: it sporadically requires to inhibit a movement upon an incoming salient stop-signal.

To study the neural basis of arm movements inhibition we combined the approach typically employed for the study of perceptual-decision making with the countermanding task, that is broadly used to evaluate response inhibition

To this aim we modified the salience of the stop-signal and we found that this modulation affected the ability to inhibit in macaque monkeys: coherently to what already observed in humans, we found that less salient stimuli deteriorate inhibitory performance. These behavioural results were subtended by neural modulations representing an inhibitory process that started later in time and showed a less steeper dynamic for stimuli difficult to be processed.

This study shows that the neural patterns observed when deciding to stop are broadly similar to the neural patterns observed when deciding to act in the literature; thus it is a first step in investigating the perceptual decision making process involved in movement inhibition.

## Introduction

Many daily decisions we make are based on the processing of sensory stimuli. For example at the traffic light, we press the accelerator pedal when the light turns from red to green. However this simply colour change detection could be less or more efficient, the same green light could be more easily detected at night than in sunshine.

Several studies have used perceptual tasks to explore decision processes. Typically, in a controlled experimental setting subjects are presented with different amount and/or quality of visual information and their performance, evaluated as selected choice or response time, reflects the amount/consistency of information progressively accumulated. Typically the larger the amount of information available (or the less is the level of uncertainty), the faster and the more accurate is the response. Those studies suggest that a decision is taken when a signal of evidence reach a threshold level in support of the action that will be made^1,2,3,4^.

One key aspect is that in perceptual tasks decisions are communicated with an action, an overt behaviour, typically a hand or an eye movement. A similar outcome is unavailable when the perceptual task triggers an action cancellation. The absence of a direct behavioural evidence is an important reason why we know much less of the processes that supports the decision to suppress a behavior. This lacking of knowledge remains despite suppression ability is of central importance in many fields of neuroscience, altered in various neuropsychiatric and neurological diseases ^5,6^.

Both frontal and parietal cortical areas of the primate brain contain neurons which activity show evidence of accumulation dynamics^3,7,8,9,10,11,12^, eventually combined with an urgency to respond process^13^. In some of these areas (e.g., the lateral intraparietal area, LIP, and the dorsal premotor cortex, PMd) sensorial information is integrated into a movement preparation activity that is later transformed in an action. Neural activities changes typically correlate with stimulus perceptual difficulty, choices and response times. For example, the easier is the stimulus to detect, or to distinguish against alternatives, the faster the accumulation of evidence and the shorter the response time. The uncertain relationship between target and action could be expressed also by tasks where multiple actions are simultaneously programmed and only one selected based after a delayed instruction cue^14^.

An open question is whether the decision to cancel an almost occurring behaviour is supported by neural dynamics similar to the ones operating in overt decisions.

To directly tackle this issue, we trained rhesus monkeys (Macaca mulatta) to perform a countermanding task, extensively employed to investigate the suppression abilities at the neural as well behavioural level ^15,16^. In this task it is required to respond to a Go stimulus, presented either to the left or to the right of the workspace in different trials (**no-signal trials**), but to halt the response when an unpredictable Stop-signal subsequently occurs after varying delays (**stop-signal delay, SSD**) in a subset of trials (**stop-signal trials**). In each stop-signal trial, one can withhold (**signal-inhibit trials**) or generate the response (**signal-respond trials**). This task allows evaluating the so-called reactive inhibition that is described by the probability to respond to the Stop-signal as function of the SSD lengths and by the duration of the stop process (**stop-signal reaction time, or SSRT**). This last one is an estimate of the time that it takes to stop the response after the stop-signal presentation^15^. The SSRT can be broadly considered the response time of the inhibitory process. As such it is supposed to be modulated by different task demands^17,18^.

In the framework of the race-model, brain regions that show neuronal activity modulations related to motor decision should be the ideal locus to investigate the dynamic of perceptual decision-making associated to the inhibitory demand.

In the frontal lobe of primates, similar characteristics can be found in the PMd known to integrate at different levels spatial information with visual information. Neural activity of this area represents various properties of the impending movement, including its direction, distance, trajectory, timing and speed ^19, 20,21,22,23,24,25,26,27, 28,29,30,31^, it signals the direction of potential and final reach choices^14^ as well the selection of specific rules^32^. Furthermore it reflects conflicts related to the dynamic competition between contemporary alternative movement choices^33,34^, and express a decision process related to the selection of actions based on sensory cues^35^ or dynamic sensory signals^34^ as well change of mind^36^. Of relevance, PMd contains neural activity that modulates according to movement inhibition^37,38^.

Here we found that, as in humans recently tested in the same task^39^, less salient stimuli deteriorate inhibitory performance and correspond to longer SSRT. These behavioural data were subtended by neural modulations in PMd, participating to action inhibition, that started later in time and showed a less steeper dynamic for stimuli difficult to be processed.

Thus actively inhibiting an action shows neural dynamics that are specular to those previously described at play for overt perceptual decision making ^2,3,6^.

## Results

### Behavioral performance and neural selection

In the stop-signal trials of the countermanding task we adopted (Figure 1A) the Stop signal corresponded to a change in colour, detectable with different level of difficulties (easy, medium, or hard), of the Go signal after a variable SSD.

**Figure 1:**
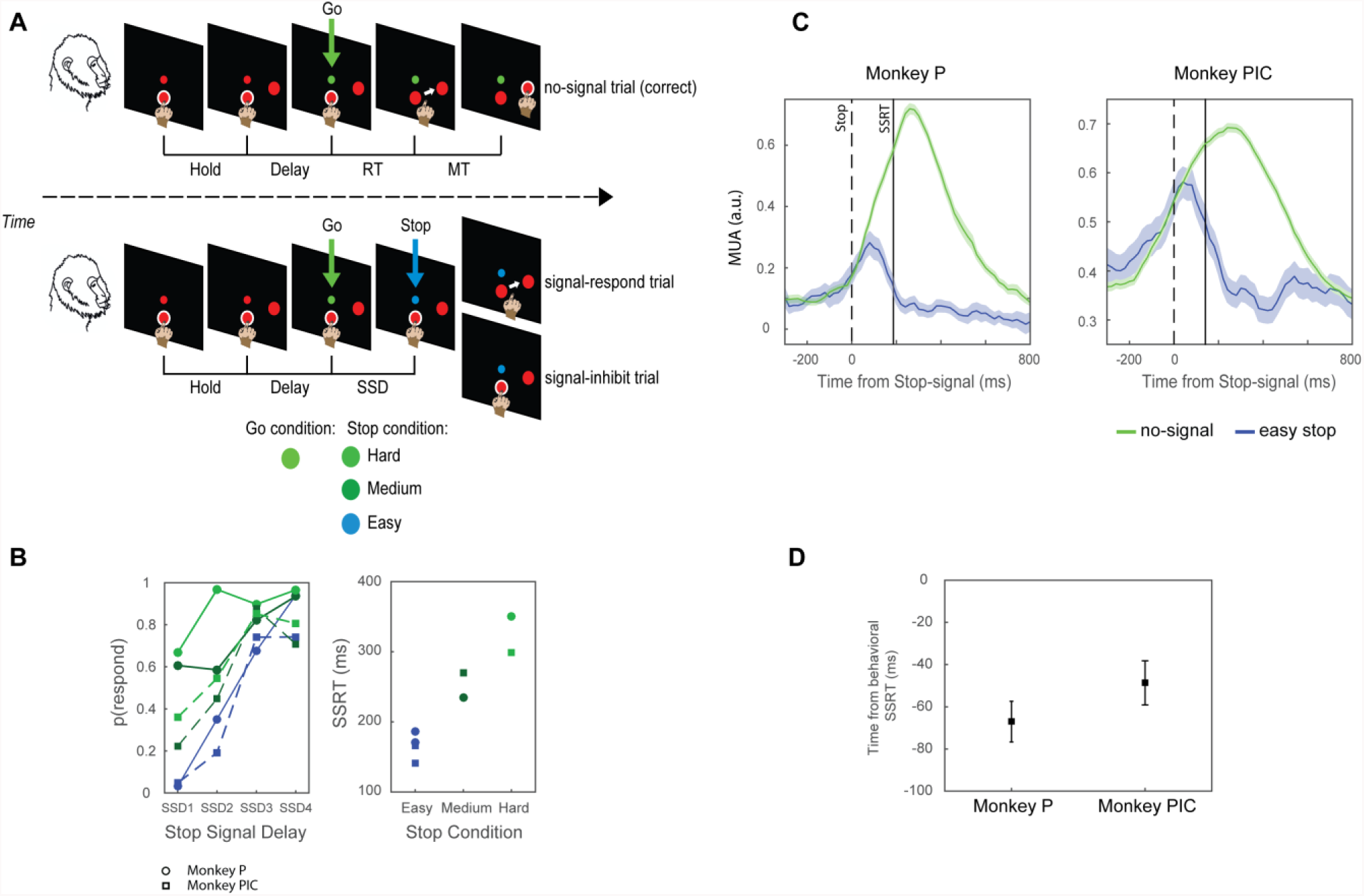
A Countermanding task (multi-stop-signal version). Monkeys touched the central target (CT) for a variable duration (Hold, depending on the session) at which expiration the peripheral target appeared and the Delay period started. Then the Go signal appeared, instructing for a reaching movement towards the peripheral target. In no-signal trials the monkeys were rewarded upon the touch of the peripheral target. In stop-signal trials the monkeys had to refrain from moving to get the reward (signal-inhibit trials); otherwise, if a movement was made, the reward was not delivered (signal-respond trials). One out of three different stop-signals could unpredictably and equally probable appears (Stop salience conditions: easy, medium, hard). B Effects of the stop-signal salience modulation on inhibitory performance: Left column: Inhibition functions obtained for the two monkeys in the fixed SSDs session; Right column: SSRT estimates separately for each monkey (see Behavioral Results for details). C Comparisons between easy signal-inhibit trials and latency-matched no-signal trials for a single movement direction across all channels that showed modulation before the end of the SSRT. Data are relative to a single recording session (Monkey P, n=; monkey PIC n= 6) D Latencies of the divergences between signal-inhibit and no-signal trials (mean±SE) relative to the finish time of the SSRT (monkey P, n=58; monkey PIC n=12).

For increasing the power and validity of the neural data analysis, we selected two behavioural sessions for each animal (Monkeys P and PIC) on the basis of the following criteria: a total of more than 1000 total trials per session; one session for the fixed-SSD procedure and one for the variable-SSD procedure.

First we will briefly describe the behavioural performance of the monkeys within the framework of the race model showing that the salience of the Stop signal was able to affect the inhibitory performance. We will then illustrate the PMd neural correlates of this behavioural effect.

Figure 1 B shows, for the sessions with fixed SSDs, that in both monkeys a lower salience of the Stop signal corresponds to a shifting of the inhibition function toward higher probability of errors for the same SSD durations. Furthermore, the values of SSRT display a similar trend with a longer duration for Stop signals hard to be detected.

For the neural analysis we will describe the changes in the modulation of multiunit activity (MUA), obtained from the unfiltered signal (see Materials and Methods) recorded from each electrode and processed as in our previous paper^40^. We initially selected data from electrodes with reaching related modulation in no-signal trials and in at least one of the two directions of movement. From this dataset we further selected those channels related to the countermanding task, i.e., we selected MUA modulations in PMd predicting whether or not a movement will be made (see Fig. 1 C for the modulation at the population level, and Supplemental figure 2 for single channel examples) by evaluating their time of divergence between no-signal and signal-inhibit trials (latency-matched only, see Material and Methods and Fig. 1D). For this last selection, we employed only signal-inhibit trials in the *easy* conditions: indeed across monkeys and sessions this condition provides reliable estimates of SSRTs (see Behavioral Results); furthermore previous neurophysiological study have employed salient stop-signals that were easy to detect to establish the role of neural activity in movement control (as such roughly corresponding to the *easy* condition^16,37,38,41,42^.

In conclusion we overall selected 58 channels for Monkey P (30 and 28, in separate sessions) and 12 for Monkey Pic (6 and 6, in separate sessions), for a depth analysis of the effect of the Stop signal detection in the changes of preparatory activity.

### The salience of the stop-signal affects PMd neural dynamics

To investigate the effect of stop signal salience on neural activity we first compared *easy*, *medium* and *hard* signal-inhibit trials aligned to the stop-signal presentation.

For both monkeys we found clear differences between the salience conditions both for single recording channels as well at the population level (see Fig. 2, A-D). When the Stop-signal was *easy* to detect, the suppression of neural activity started at about 130 ms after Stop-signal presentation; differently it was delayed for the *medium* and *hard* conditions (up to about 220ms). Furthermore the neural dynamics appear to be different: in the *easy* conditions compared to the others a steeper suppression occurred.

**Figure 2.**
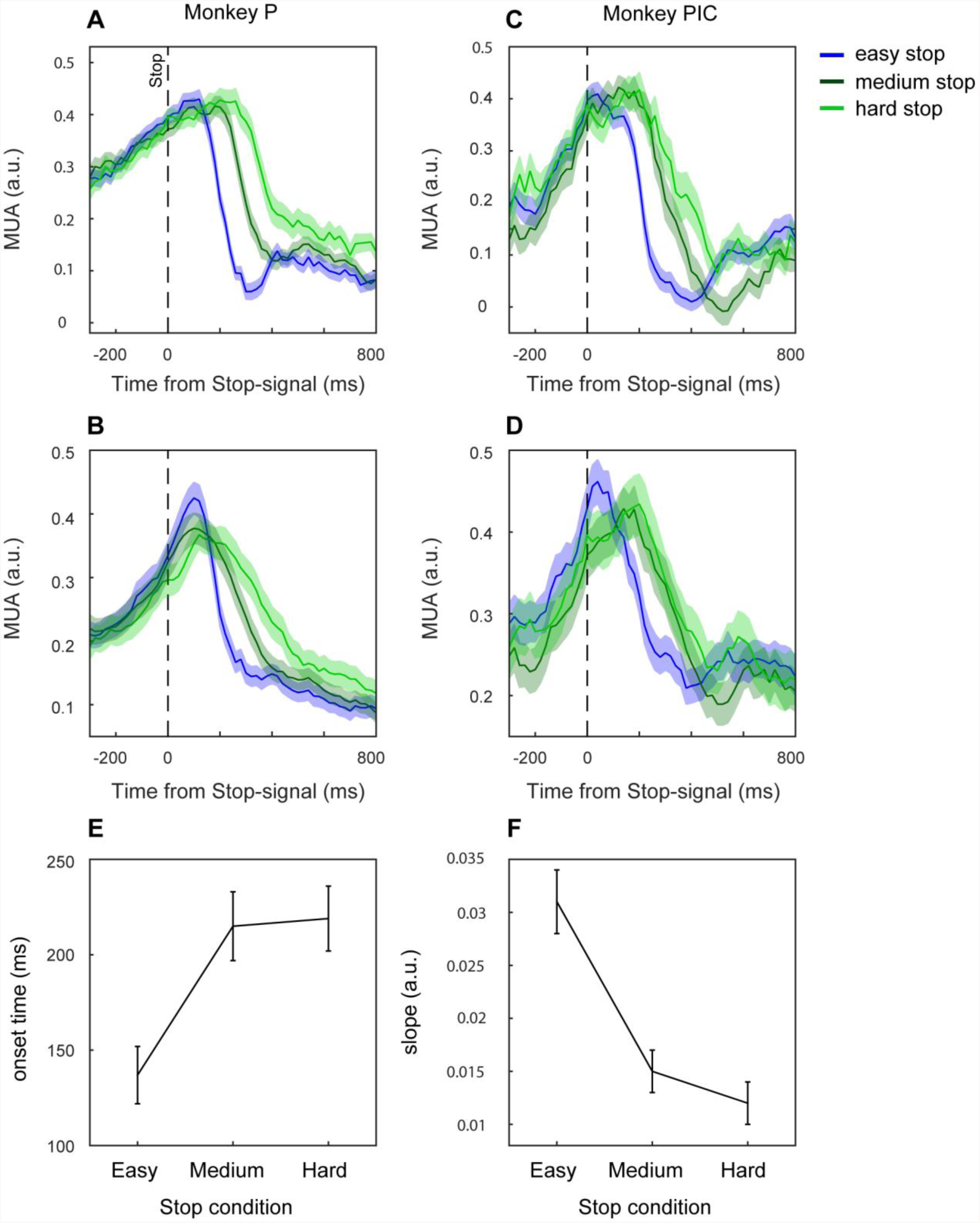
Effect of salience conditions on MUA. Neural activity is aligned to the Stop signal and it is represented for a single channel (A and C) and at the population level for a single session (B and D) separately for each monkey (A,B: monkey P, for the population n=28; C,D: monkey PIC, for the population n=6). Onset times (E) and slopes (F) of the neural modulations across all sessions and monkeys (mean±SE).

Statistical testing supported the phenomenological pattern of neural activities in Figure 2: two separate ANOVAs, one for the latency of the onset of suppression relative to the Stop-signal (onset times), and the other for the slopes, with factors monkeys and stop-signal conditions, were done. For the slope analysis we found that the slopes changed depending on the salience (F(2, 116)=32.4, p=.00000): the *easy* condition showed a steeper slope (mean= .031; 95 CI= 0.024, 0.037) compared to the *medium* and the *hard* conditions (*medium*: mean=0.015; 95 CI= 0.011, 0.019; *hard*: mean=0.012; 95 CI = 0.008, 0.016; Newman-Keuls post-hoc test: MSE= .00012, df = 116, *easy* vs *medium* p=.0001; *easy* vs *hard* p=.0001); however *medium* and *hard* were not different between each other (p=0.45)).

The analysis of the onset times showed that salience conditions affected the inhibition process (F(2, 114)=12.231, p=.00002; Newman-Keuls post-hoc, MSE= .00680, df = 114): specifically in *easy* trials MUAs was suppressed earlier (mean=136, 95 CI= 105, 167) than in *medium* and *hard* conditions (medium: mean= 214, 95 CI=178, 251; p=0001; hard: mean = 218, 95 CI=183 254, p=.0001, respectively); however no differences were detected between *medium* and *hard* conditions (p=0.4).

Thus in general we found that the higher the salience of the Stop-signal, the earlier the modulations of neural activity starts and the stronger (steeper) the modulation observed.

### Interference of the stop process on the go process in signal-respond trials

In the behavioural analysis we found that the independence assumption was confirmed for the easy conditions but not always for the medium and hard. One possible explanation is that in the other conditions, the stop process interacted with the go-process in signal-respond trials, lengthening its duration. Another, alternative explanation, is that the higher length of the stop process in medium and hard condition give the possibility to have longer RT in signal-stop trials, thus making more difficult to find difference between no-signal and signal-respond trials. We found already evidences, at the neural level, that the inhibition process is longer in the medium and hard condition. Indeed it starts late and its dynamic shows a slower suppressive modulation. However this effect could be combined with an interference effect, or, in other terms, both altered neural dynamic and interference can participate in producing longer signal-respond RT. To evaluate whether some interference was active in signal respond trials, we compared the activity of signal-respond trials, separately for each conditions, to latency matched no-signal trials. If an inhibitory process had been active in signal-respond trials, the corresponding neural activity should have been lower than the selected no-signal control trials, where no inhibitory process was active. To this aim we considered as latency matched the no-signal trials with RT shorter than the average SSD of all signal-respond trials plus the SSRT estimated for each session in the easy condition. We then calculated the average MUA in the −180 −80 ms interval aligned to the time of detach. We considered the same interval also for all the signal respond-trials in each condition. We found a three way interaction between the factors considered (monkeys, sessions, and type of trials: F(3, 198)=6.5, p=.00034) and we ran Dunnet post-hoc tests).

Here we report the data separately for each monkey and session: Monkey P, fixed SSDS, mean (SE): no-signal 0.46(.06); signal-respond: *easy* (0.46(0.06), *medium* (0.46(0.06); *hard* 0.51(0.06)), all p’s >0.63; Monkey P, staircase: no-signal 0.6(0.06); signal-respond *easy* (0.59(0.06)), *medium* (0.6(0.06)), *hard* (0.6(0.06)) (all p’s >.8); Monkey PIC, Fixed SSDs: no-signal (0.74(0.13)), *easy* (0.73(0.14), *medium* (0.74(0.14)), *hard* (0.73(0.14))) p’s > 0.62; Monkey PIC, Staircase: no-signal (0.37(0.14)), *easy* (0.38(0.14), *medium* (0.39(0.14)), *hard* (0.37 (0.14) (all p’s>0.95); (MSE =.111, df = 66.297. These data show that failure of independence assumption for signal-respond trials was not supported by a suppressed activation of neural activity. To further strengthen our observation we performed the same comparison by considering all no-signal trials. Qualitatively identical results were obtained (three way interaction analysed by means of Dunnet post-hoc tests: F(3, 198)=3.9, p=.009, all p’s of comparisons > 0.8).

Thus we can exclude that, at least at the level PMd, interference between stop and go processes cannot explain the failure of independence assumption and the lengthening of signal-respond trials (see Figure 3)

**Figure 3.**
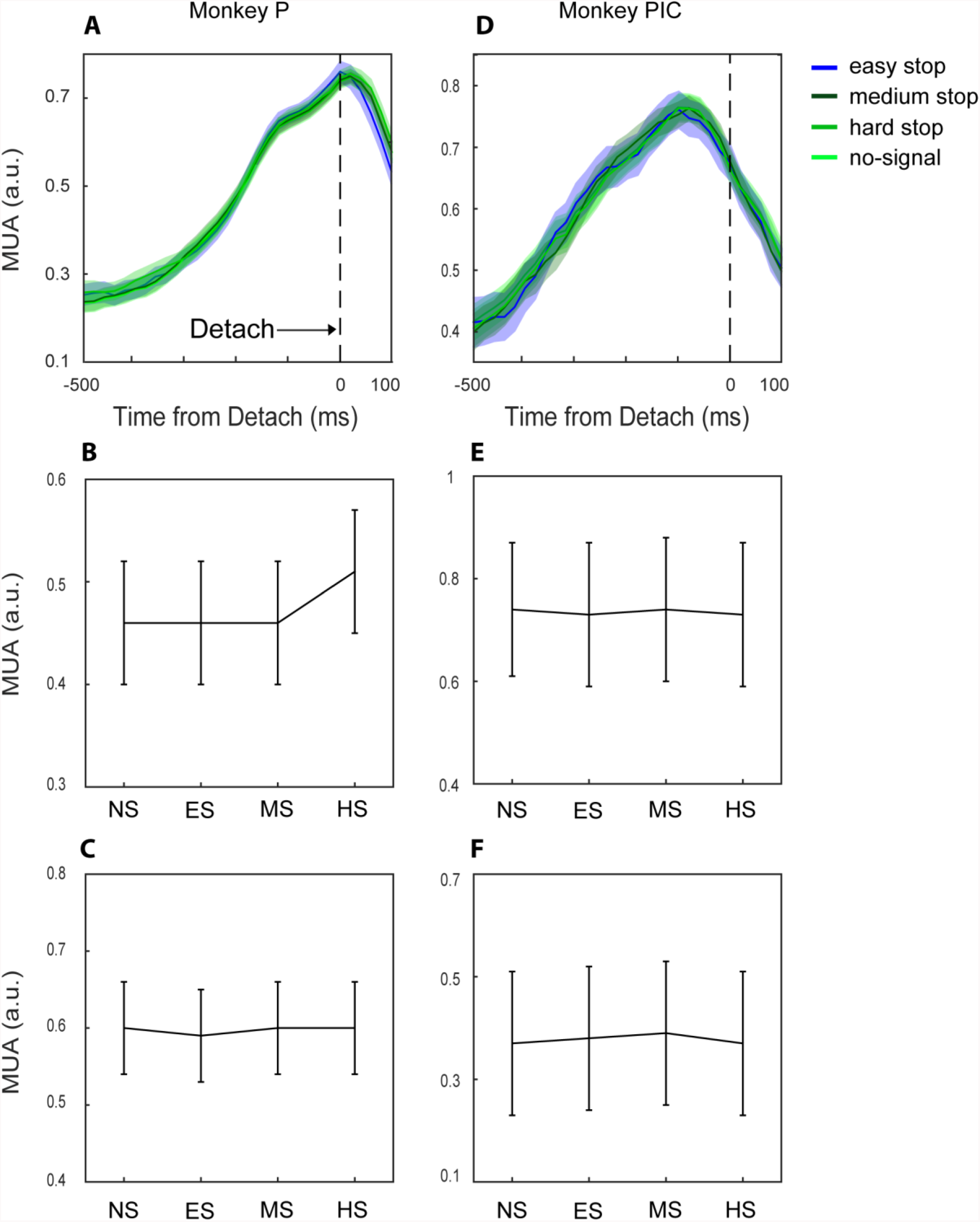
Comparison between no-signal and signal-respond trials. First row: population average comparison for a single session (A, monkey P, n=28; B, monkey PIC, n=6). Second row (C,G): comparison between average activities before movement onset across monkeys for fixed SSD sessions (mean±SE); Third row (D,H): comparison as above for staircase sessions across monkeys (mean±SE).

## Discussion

Aim of the present study was to investigate the neural activity in PMd related to perceptual decision during inhibitory process. We used a countermanding task in which the salience of the Stop-signal was modulated: we found that the higher the salience the higher the ability to stop. This behavioural performance is well predicted by the neural activity. Following a clear Stop-signal, we observed a modulation of neural activation driven by the Stop-signal that started earlier and showed a steeper slope, suggesting a “stronger” decision process, compared to Stop-signals that were less salient.

This is, to the best of our knowledge, the first behavioural and neurophysiological study that explicitly combines the perceptual modulation of the stop command in the context of the countermanding task.

The temporal patterns that we observed are reminiscent of studies of the neural mechanism underlying decision involving perceptual processing and movement selection. For example studies using RDM (Random-Dot Motion^43^) require making saccades towards the target corresponding to the direction by which the highest fraction of dots are coherently moving. Increasing the fraction of dots that moves coherently, and thus the strength of the stimulus, make the task easier. Indeed accuracy increases and RT decreases for strong stimuli. Neural activity in parietal areas (e.g., LIP) reflects the decision process about where and when to move on the basis of the fraction of coherent motion detected: the stronger the stimulus the steeper the accumulation of evidence and the shorter the RT^2^. There are similarities between our task and the pattern described above. We do not have a random dot Stop-signal, but the modulation of the visual salience clearly affects the inhibitory performance. If we consider the decrease of activity that we observed as the reflection of an active process against the movement generation, we can see that its temporal evolution parallels the signal that reflects accumulation of evidence typically described.

In the last years different groups have addressed the role of PMd in somatomotor decision by employing tasks in which the sensorial instruction required some form of discrimination (establishing which amount of two colours was more present ^33, 35^) or could continuously change ^13^. In all these studies neural activity has been show to reflect process related to the decision formation either with or without accumulation of evidence^34^. In a study aimed at investigate the cortical laminar differences of processing in a decision making task, it has been found that a subpopulation of neurons prevalent in the superficial layers showed a build-up of neural activity that was steeper for easy and faster responses, thus reflecting the decision process^35^. Thus these studies give supports to the hypothesis that the pattern we observed is a reflection of a decision-to-stop process that is modulated by the visual salience.

Previous neurophysiological investigation on movement inhibition in primates, by means of the countermanding task, recorded neural activity from FEF and SC, and employed an easy to detect Stop-signal^16,41^. The same type of Stop-signal has been obtained in studies aimed at investigating the neural correlates of arm movement inhibition^37,42^. At the same time behavioural studies have shown that sensory features of the Stop-signal or presence of distractors can affect inhibitory performance^39,44,45,46,47,48,49^. Furthermore simulation studies have investigated the role of the detection process in permitting the inhibition of saccades^50^ and how response inhibition can result from blocking the input to the Go process thus avoiding its growth to generate the movement. However so far no direct neurophysiological investigation has been performed. We think that our work it is seminal in showing the neural consequences of the modulation of the perceptual processing during movement inhibition, and it will provide new insight for the refinement of neural inspired model of movement control^50,51,52,53^.

Another important contribution of our results it is to confirm the role of PMd as a site of movement suppression. First we have shown that MUAs conveys a signal sufficient to predict whether or not a movement will be inhibited, as observed for single neurons^37^, and as suggested by movement planning dynamics also in other tasks^40^. MUA closely approximate the firing activity of small population of neurons nearby the electrode.

Note that so far, at the high temporal and spatial definition of neurophysiology, this is the only cortical area that shows such a strong relationship with arm movement suppression.

Together with PMd, also SMA has shown field potential modulations in temporal relation with movement inhibition in the countermanding task. However it must be considered that these field potentials can be related to widespread phenomena that probably reflect an incoming signal from other regions (like prefrontal cortex or basal ganglia), as also observed in different contexts^54,55,56,57,58,59^. Whether or not this signal will be used to stop the movement, or to regulate other aspects of the behaviour related to the task, can probably be established by considering the MUA or single cells firing rates^60^. We propose that PMd will provide the output to other cortical or subcortical structures as suggested in other neurophysiological and neuroanatomical studies^31,61,62,63,64,65,66,67,68^.

The data we presented further supports the role of PMd in movement inhibition because the neural modulation in the different conditions covaried with the behavioural performance. Although our data are correlational in nature, support to a role of PMd comes also from TMS studies^69,70,71,72,73^ as well lesions in both monkeys and humans ^74,75^.

A role of PMd in humans has been also confirmed by employing electrocorticography^40^. Such a support has partially been provided by fMRI studies^76,77^: probably the nature of the signal, and the wide temporal definition makes difficult to extract a signal related to the neural dynamics of this region.

Another aspect of this study is that, from the behavioural point of view, performance in monkeys is very close to the performance observed in humans^39^ suggesting that the same neural processes can be at play. This possibility is of particular importance when considering the application of basic neurophysiological knowledge to help clinical investigations. We still have to deeply investigate the neural processes that support the decision to suppress a behaviour: this lacking of knowledge occurs despite suppression ability is of central importance in many fields of neuroscience, because it seems to be damaged in various neuropsychiatric as well neurological diseases^5,78^. For example detriment in inhibition performance is often observed in Attention Deficit Hyperactivity Disorder (ADHD) persons performing the stop-task^47,79,80,81^. However it is not clear which are the specific components that are affected ^82,83^. Considering the present results, diverse possible mechanism could contribute in reducing the ability to control the movement: a diminished triggering of the stop-response, the delayed onset of stop implementation or the slow rate of implementation of the same process.

Neurophysiological inspired modelling could help in predicting specific behavioural performance on the basis of malfunctioning of different components, thus helping in understanding the neurocognitive basis of neuropsychiatric and neurological disorders.

## Materials and Methods

### Subjects

Two male rhesus macaque monkeys (Macaca Mulatta, Mon P and PIC) weighting 9 and 13 kg were employed for this study. Monkeys were pair-housed with cage enrichment. They were daily fed with standard primate chow supplemented with nuts and fresh fruits if necessary. The monkeys received their daily water supply during the experiments. All experimental procedures, animal care, housing, and surgical procedures conformed to European (Directive 86/609/ECC and 2010/63/UE) and Italian (D.L. 116/92 and D.L. 26/2014) laws on the use of nonhuman primates in scientific research and were approved by the Italian Ministry of Health.

### Animal preparation

In both monkeys a 96 channels Utah arrays (BlackRock Microsystem, USA) was implanted in the PMd using anatomical landmarks (arcuate sulcus - AS - and pre-central dimple - pCD) after dura aperture. All the surgeries were performed under sterile conditions and veterinary supervision. Antibiotics and analgesics were administered postoperatively. Anaesthesia was induced with ketamine (Imalgene, 10 mg kg^−1^ i.m.) and medetomidine hydrochloride (Domitor, 0.04 mg kg^−1^ i.m. or s.c.) and maintained by inhalation isoflurane (0.5–4%) in oxygen (5 l min−1). Antibiotics were administered prophylactically during surgery and postoperatively for at least 1 week. Postoperative analgesics were given at least twice per day. Recordings started well after recovery from surgery (minimum after 10 weeks). A head holding device was implanted in monkeys PIC before training, while in monkey P the head holder was implanted simultaneously with the array (see below).

### Apparatus and tasks

Experiments were performed in a darkened acoustic insulated room Monkeys seated in front of a black isoluminant background (<0.1 cd/m2) of a 17-inch touchscreen monitor (LCD, 800 × 600 resolution). A non-commercial software package, CORTEX (ftp://helix.nih.gov/lsn/matoff/), was used to control stimuli presentation and behavioural responses.

Figure 1 shows the schema of the general task employed, which is a modified version of the countermanding task, characterized by the presentation of two types of trials, randomly intermixed: no-signal trials and stop-signal trials (depending on the session the probability of stop-signal trials was between 27 and 36%). This task it is characterized by the presence of different Stop-signals.

Each trial started with the appearance of a central target (CT) (red circle, diameter 1.9 cm) and a Cue signal (red circle, diameter 0.7 cm) slightly above (3 cm) the CT at the center of the screen. Monkeys had to touch the CT and maintain their finger on it.

After a variable delay a peripheral target (PT) (red circle, diameter 1.9 cm) appeared randomly in one out of two possible locations (i.e., 7 cm at the right or left of the screen vertical midline for Monkey P only at the right for monkey PIC).

In no-signal trials, after a fore-period delay (ranging from 120 ms to 1200 ms), a Go stimulus, consisting of a green circle, appeared, replacing the Cue (no-signal condition; stimulus circle 0.7 cm RGB: [0 250 0]). The Go stimulus instructed the subjects to reach the peripheral target as fast as possible and to hold the new position for a variable time (600–800 ms), until the end of the trial. Reaction times (RTs) were defined as the time between the Go stimulus presentation and the hand movement onset towards the PT.

In stop-signal trials, the sequence of events was exactly the same until the Stop-signal (circle, 0.7 cm) replaced, after a variable delay (stop-signal delay or SSD), the Go stimulus. In these trials, the simple detaching of the hand after the Go stimulus presentation corresponded to a wrong response. Conversely, a hand kept on the CT until the end of the trial (800–1000 ms) corresponded to a correct response (signal-inhibit trial; Fig. 1, lower right panel).

For both no-signal and stop-signal trials movements performed before the Go stimulus aborted the trial, and trials were considered as not engaged, and excluded from further analysis. For correct no-signal trials and signal-inhibit trials monkeys experienced a brief sound accompanied by the delivery of the juice reward. In signal-respond trials, neither sound nor reward was delivered, and the screen became blank.

The inter-trial interval was set at 800 ms. Three different types of equiprobable Stop-signals, distinguishable in colour and brightness, could follow the Go stimulus, depending on the stop-signal condition. These Stop-signals were classified as easy, medium and hard (no-signal condition; stimulus circle 0.7 cm RGB: [0 250 0]; SS_easy [0 0 188]; SS_medium [0 160 0]; SS_hard [0 210 0]). We made the Stop-stimulus progressively closer to the Go stimulus in terms of appearance (see Fig.1).

Different procedures were adopted to establish the duration of the SSDs: tracking and fixed. By using the tracking procedure we controlled the duration of the SSDs on the basis of the performance in the last stop-signal trial: if the monkey succeeded in withholding the response, the SSD increased by one step in the subsequent stop-signal trial. Conversely, if the subject failed, the SSD decreased by one step. Each stop-signal condition had its own independent staircase procedure.

### Behavioural Analysis

The performance in the countermanding task is accounted by the race model: in stop trials two stochastic processes race toward a threshold: the GO process, and the STOP process. The result of this race, either movement generation in signal-respond trials or movement suppression in signal-inhibit trials, will depend on which of these process first will surpass its own threshold. In signal-inhibit trials the STOP process wins over the GO process and vice versa. By making the SSDs unpredictable and variable the output of the race is affected: the longer the SSD, the higher the probability to facilitate the GO process. To make the employment of the race model fruitful to study response inhibition, a central assumption must be satisfied: the GO process in the stop-signal trials must be the same as in the no-signal trials (independence assumption ^15,84^). To broadly validate this assumption signal-respond trials RTs must be shorter than the no-signal trials. All the sessions included in this study respected this prediction for at least one stop-signal condition, typically the easy condition (see Results). Once the independence assumption has been validated, Stop-signal Reaction Time (SSRT) can be estimated. This is done by extracting three main variables: the reaction time (RT) distributions of no-signal trials and signal-respond trials, and the probability to respond (p(R)) by error to the Stop-signal. Following recent recommendations ^85^, we first confirmed that the independent assumption was respected and then we proceeded in calculating the SSRT by using the “integration method” ^84^. Because our behavioural analysis were performed at the single session level, we adopted a comparison criteria between average values because in these conditions the employment of a statistical test can be a too strict criterion^85^. However, in the easy conditions, for 3 out of 4 session the difference was also statistically confirmed (ranksum test, see Appendix of Results).

The method to estimate SSRT assumes that the finishing time of the stop-signal process corresponds to the *n*th no-signal RT, where *n* results from the multiplication of the ordered no-signal RT distribution by the overall probability of responding, *p*(respond|signal) when using the tracking procedure. SSRT can then be calculated by subtracting the mean SSD from the *n*th no-signal RT. When the fixed procedure was employed, we applied the same method at each SSD (provided the *p*(respond|signal) was above 0 and below 1). We then averaged SSRTs to obtain a single value.

### Neural recordings and analysis

Neural activities were recorded from 96 channels Utah arrays (BlackRock Microsystem, USA) by using specific software (Tucker Davies Technologies, Unfiltered raw signal, Sampling rate 24.4Khz). Array data in this paper come from two recording sessions for both monkey P (2 months interval between sessions) and monkey PIC (6 months interval). The Multi Unit Activity (MUA) that we extracted is a good approximation of the average firing rate as described in details in 40,86In this study, we smoothed the signal obtained by using a moving average sliding window (±40 ms sliding window (20 ms step). We selected channels displaying a reaching-related MUA modulation for at least one movement direction i.e., a significant difference between the activity preceding the movement onset (from − 250 to −50 ms before detach) and the baseline (from 0 to 100 ms following the touch of the CT). This type of modulation is an index that neural activity recorded from that channel is potentially involved in movement preparation and control. For the main analysis reported here where focused on channels which MUA was modulated before the end of SSRT as done in other works in primates on the same topic^16,41,42,37,38^.

To compare neural activities between signal-inhibit and no-signal trials, we considered a subsample of the last group of trials: the so called latency-matched trials. Those are trials which RTs where longer than the specific SSD (or average SSD) plus the corresponding behavioural SSRT.

When no-signal were compared to signal-respond, the latency matching was performed by considering the no-signal trials with RTs longer than 100 ms and shorter than SSD+SSRT. Latency-matched are trials that have the same level of movement preparation of the target trials for the comparison (either signal-inhibit or signal-respond).

To select neural responses involved in movement suppression we established whether MUA modulation could predict movement inhibition by estimating when the divergence between signal-inhibit and latency matched started. To gain more power, we aligned the signal-inhibit trials to the stop signal presentation, while the latency matched trials were realigned by considering the average SSD employed in the signal-inhibit trials. We calculated a differential MUA function by subtracting the average MUA in signal-inhibit in the easy condition from the latency matched no signal trials. We defined as time of the divergence the bin at which the differential MUA exceeded by two SD the mean difference in the 200 ms preceding the stop signal presentation and stayed above this limit for at least 6 bins (120ms).

From the values obtained we subtracted the corresponding SSRT and obtained a negative value, indicating the time lag between neural and behavioural estimate of inhibition. Negative values indicates that the neural modulation occurs before the behaviour.

To evaluate the latency of the MUA modulation following the Stop signal, we identified the time, from the stop presentation, in which MUA began to show a decreasing trend that went on for at least 100 consecutive ms. This time was defined as onset time. Starting for the onset time we considered the following 200 ms and we ran a robust regression function (Matlab robustfit) to extract the slope of the MUA suppression pattern.

This analysis was repeated for each selected channel and stop condition across all the recording sessions for both monkeys. The datasets generated and/or analyzed for the current study are available from the corresponding author on reasonable request.

## Acknowledgments

The authors thanks Maurizio Mattia for providing tools for the extraction and preprocessing of the neural signal.

## Appendix

Detailed behavioural results session

**Table 1:** for different monkeys and sessions the following data are reported: overall number of trials (n trials); percentage of stop-signals presented (% Stop); Mean and SD of Reaction Times (RTs) for no-signal, signal respond easy, medium, hard trials (s-r Easy, s-Medium, s-r Hard) together with the p value from the comparison between no-signal and the target signal-respond trials groups. Stop signal Reaction Times (SSRT) are reported only for comparisons for which signal-respond trials RTs were not nominally higher than no-signal RTs.

| Monkeys Session | n trials | % Stop | no-signal RT Mean(SD) | s-r Easy RT Mean(SD); p | s-r Medium RT Mean(SD); p | s-r Hard RT Mean(SD); p | SSRT easy | SSRT medium | SSRT hard |
| --- | --- | --- | --- | --- | --- | --- | --- | --- | --- |
| P(fixed) | 1228 | 33 | 637.9 (126.1) | 589(114); p=.001 | 642(136); p=.9 | 638(112); p=.9 | 171(84); |  |  |
| P(staircase) | 1399 | 36 | 565(104) | 526(82); p<.001 | 548(95); p=.2 | 562(122); p=.4 | 186 | 235 | 350 |
| Pic(fixed) | 1131 | 27 | 627(177) | 518(154);p<.0001 | 569(120); p=.0042 | 569(158); p=.0042 | 166(68); | 270(122); | 299(114) |
| Pic (staircase) | 1517 | 30 | 387(126) | 357(133); p<.3 | 397(158); p=.5 | 412(186); p=.5 | 141 |  | 2 |

